# Superior temporal sulcus hypoperfusion in children with autism spectrum disorder: an arterial spin-labeling magnetic resonance study

**DOI:** 10.1101/771584

**Authors:** Ana Saitovitch, Elza Rechtman, Hervé Lemaitre, Jean-Marc Tacchella, Alice Vinçon-Leite, Elise Douard, Raphael Calmon, David Grévent, Anne Philippe, Nadia Chabane, Francis Brunelle, Nathalie Boddaert, Monica Zilbovicius

**Affiliations:** INSERM U1000, Department of Pediatric Radiology, Hôpital Necker Enfants Malades, AP-HP, University René Descartes, Pres Sorbonne Paris Cité, Institut Imagine and UMR 1163, Paris, France; Faculté de Médecine, Université Paris Sud, Saclay, France; UMR 1163, Paris Descartes University, Sorbonne Paris Cité, Institute IMAGINE, Paris, France; Centre Cantonal Autisme CHUV, Centre hospitalo universitaire Lausanne, Lausanne, Suisse

**Author notes:** both authors contributed equally for this work (Ana Saitovitch, Elza Rechtman). Corresponding author’s contact information: Ana Saitovitch.

## Abstract

Advances in neuroimaging techniques have significantly improved our understanding of the neural basis of autism spectrum disorder (ASD). Several attempts have been made to label the main neuroimaging phenotype of ASD, mostly by anatomical and functional activation studies, but none of the frameworks have been without controversy. Over the past decade, a renewed interest for rest brain functioning has emerged in the scientific community, reflected on a large number of resting state fMRI (rs-fMRI) studies, but results remain heterogeneous. It is possible today to investigate rest brain functioning by measuring rest cerebral blood flow (CBF) with MRI using arterial spin labeling (ASL). Here, we investigated rest CBF abnormalities using non-invasive ASL-MRI in 18 children with ASD without cognitive delay (10.4 ± 2.8 y) and 30 typically developing children (10.6 ± 3.0 y). Following quality control, images from a final sample of 12 children with ASD (11.2 ± 2.9 y) and 28 typically developing children (10.1 ± 2.5 y) were analyzed. Whole brain voxel-by-voxel analysis showed significant rest CBF decrease in temporal regions, mainly in the superior temporal sulcus (STS), in children with ASD. This hypoperfusion was individually detected in 83% of children with ASD. Finally, negative correlation was observed between ASD severity scores and rest CBF in the right posterior STS. Strikingly, despite the small sample studied here, our results are extremely similar to previous PET and SPECT findings describing decreased rest CBF in the same superior temporal regions at group and individual levels, as well as correlation with symptoms severity. The congruence between these results, with different methods and in different ASD profiles, reinforce the strength of rest functional abnormalities within these superior temporal regions in ASD and strongly indicates it might be a core characteristic of the disorder. Identifying a core dysfunctional region in ASD bears direct implications to the development of novel therapeutic interventions, such as transcranial magnetic stimulation. In addition, if confirmed in a larger sample, rest temporal hypoperfusion could become a reliable brain imaging biomarker in ASD.

## Introduction

Autism spectrum disorders (ASD) are neurodevelopmental disorders characterized by social communication deficits and restricted repetitive behaviors [1]. Major advances in neuroimaging techniques in the past few decades have allowed a wide investigation of the neural basis of ASD and greatly enriched our understanding of neuropathological features in this disorder. Brain imaging studies in ASD constitute nowadays a very large literature of more than 3000 publications (PubMed).

Even though it is generally recognized that ASD is related to structural [2–4] and functional [5–7] network abnormalities, description of its extent and overall pattern has been rather heterogeneous. Indeed, while several attempts have been made to label the main neuroimaging phenotype of ASD, none of the frameworks have been without controversy. Inconsistency may result from methodological issues such as differences in subject inclusion criteria, variability in image processing and analysis methodology [8, 9].

From a functional perspective, the first results describing abnormalities in ASD came from positron emission tomography (PET) and single photon emission computed tomography (SPECT) investigations on rest cerebral blood flow (CBF). Results from these studies showed rest hypoperfusion located in the temporal regions in children with ASD compared to control group [10, 11]. In addition, rest hypoperfusion in the temporal regions were correlated to the severity of ASD symptoms [12]. Despite these interesting findings, PET and SPECT methods presented important methodological limitations, predominantly due to the use of radiotracer injection. Thus, with the arrival of functional magnetic resonance imaging (fMRI) technique using activation paradigms, rest CBF measures by nuclear imaging methods was progressively abandoned and rest functional abnormalities in ASD have not been described since then. From the late 90’s, brain function in ASD has been mostly investigated using activation fMRI paradigms, with focus on different brain systems. Activation abnormalities have been largely described but, as intrinsic to the method, depend on the type of tasks executed in the scanner.

Over the past decade, a renewed interest for rest brain functioning has emerged in the scientific community. Resting state fMRI (rs-fMRI), a technique based on the intrinsic synchronous activity that occurs in distant regions of the brain at rest [13], has been used for this type of investigation. Resting-state fMRI studies conducted in ASD have suggest diminished connectivity, particularly between nodes of the default mode network (DMN) [14–16] and between the anterior and posterior insula and a number of social processing brain regions in ASDs [17–19] but also increased connectivity, for instance between multiple striatal regions and striatal hyperconnectivity with the pons [20]. Indeed, general over-connectivity, but also under-connectivity, and/or a combination of both have been reported in ASD, and hallmark connectivity patterns are still unclear [21]. Importantly, results should be interpreted with caution since it has been shown that motion can generate false positives where no true differences exist [22] and even small amounts of head motion during scanning have large effects on functional connectivity measures [23]. Therefore, head motion must be taken into account in MRI studies. Moreover, rs-fMRI analyses require prior hypotheses on the connected regions preventing fortuity findings, as it’s possible when using a whole brain approach.

If we consider investigations of neural basis in ASD, rest functional studies are highly pertinent since they do not require subject’s active cooperation and can therefore include the whole ASD spectrum. The emergence of a novel MRI technique offers new opportunities for assessing brain functioning at rest based on rest CBF measures. Arterial spin labeling (ASL) is a non-ionizing and completely non-invasive MRI method that uses magnetically labeled blood water as an endogenous tracer for quantification of brain perfusion, providing rest CBF measurements. In this technique, the diffusible tracer is magnetically labeled arterial blood water protons, produced by saturating or inverting the longitudinal component of the MR signal using a radiofrequency pulse If all the label arrives at the capillary bed or tissue at the time of imaging, this results in a T1-weighted signal reduction proportional to CBF, called the tagged image, which is compared to a control image, in which the blood water molecules are not perturbed [24]. ASL-MRI provides an absolute, quantifiable CBF measurement in physiological units of ml/g/min on a voxel-by-voxel basis. Finally, short acquisition time and absence of radioisotopes injections, make MRI-ASL very suitable for brain functioning at rest studies, in typical and abnormal brain development in pediatric populations.

In this study, we aimed to use this novel non-invasive method, ASL-MRI, to investigate brain functioning at rest, by measuring rest CBF, in children with ASD compared to typically developing children. In addition, we aimed to investigate whether putative rest CBF abnormalities in children with ASD measured with ASL-MRI would be correlated with the severity of autistic symptoms.

## Materials and Methods

### Participants

Eighteen children with ASD (mean age = 10.4 ± 2.8 years; four girls) and thirty typically developing controls (mean age = 10.6 ± 3.0 years; ten girls) participated in this study. Following quality control, images from six children with ASD and two typically developing controls were excluded from the study. Our final sample was then composed of twelve children with ASD (mean age = 11.2 ± 2.9 years; three girls) and twenty-eight typically developing children (mean age = 10.1 ± 2.5 years; nine girls) (Table 1). Children were diagnosed according to the DSM-IV and ADI-R criteria for ASD. All children with ASD were verbal, had an IQ score within the normal range, attended mainstream schools, had no associated neurological conditions and were free of medication for at least 1 month before imaging. All typically developing children had no psychiatric, neurological or general health problems, as well as no learning disabilities and had a normal schooling. All children from both groups had a normal anatomical MRI validated by a specialist neuro-radiologist. Written informed consent to participate in this study was obtained from each participant’s parent or legal guardian and adhere to the principles of the Helsinki Declaration. The study was approved by the Ethical Committee of Necker Hospital, Paris, France.

**Table 1.** Final Sample Characteristics

| Characteristic | Mean<br>(SD)<br>[Range] |  | P Value |
| --- | --- | --- | --- |
|  | ASD<br>(n=12) | TD<br>(n=28) | ASD vs TD<br>(n=40) |
| Sex, F/M | 3/9 | 9/19 | 0.94 |
| Age, yrs. | 11.2<br>(3.0)<br>[7-16] | 10.1<br>(2.5)<br>[6-17] | 0.256 |
| IQ | 103<br>(24.7)<br>[86-143] | 111<br>(11.1)<br>[89-140] | 0.282 |
| ADI-R score<br>(B + Cnv + D) | 42.4<br>(8.4)<br>[27-54] | NA | NA |

Clinical severity of ASD symptoms was assessed using the Autism Diagnostic Interview - Revised (ADI-R) algorithm. This semi structured investigator-based interview is based on scores of three functional domains with diagnostic thresholds. Each score contains four items. Score B quantifies impairments in reciprocal social interactions; score C quantifies impairment in communication with a verbal sub-score (Cv) and a nonverbal sub-core (CnV); score D quantifies restricted, repetitive, and stereotyped patterns of behavior and interests [25]. Because each item is scored from zero (the symptom is absent or cannot be assessed) to three (the symptom is strongly present), a child who does not speak at all (71% in this group) scores 0 on the CV score and earns a lower global ADI-R score (less severe autism) than a child able to speak with some verbal communication abnormalities. Therefore, our correlation analysis was based on a modified global ADI-R score which excluded the CV subscore [12].

### MRI scans

Perfusion images measuring cerebral blood flow (CBF) at rest were acquired for all participants using a 3D pulsed continuous arterial spin labelling (ASL) sequence with spiral filling of the K space (TR/TE: 4428/10.5 ms, Post Labeling Delay: 1025 ms, flip angle: 155°, matrix size: 128 x 128, field of view: 24 x 24 cm, with 34 axial slices at a thickness of 4 mm). CBF quantification from ASL tagged and control images was performed and warrant by General Electric 3D ASL software [26]. In addition, all participants underwent a 3D T1–weighted FSPGR sequence (TR/TE: 16.4/7.2 ms, flip angle 13°, matrix size: 512 x 512, field of view: 22 x 22 cm, with 228 axial slices at a thickness of 0.6 mm). All scans were acquired with a 1.5 Tesla (Signa General Electric) scanner at the Necker Hospital, Paris, France.

### Scans quality control

We performed a strict visual quality check on all individual T1 and ASL screening for artifacts such as motion, aliasing, ghosting, spikes, low signal to noise ratio. We excluded any images from both groups compromised by these types of artifacts (6 images from the ASD group and 2 images from the control group). Thus, all analyses presented in the following sections include the remaining 40 scans only.

### ASL-MRI pre-processing

Structural T1-weighted and ASL images were analyzed using SPM8 (http://www.fil.ion.ucl.ac.uk/spm) implemented in Matlab (Mathworks Inc., Sherborn, MA, USA). Briefly, structural T1 images were segmented into grey matter, white matter and cerebrospinal fluid and spatially normalized in the MNI space using the cat12 toolbox (http://dbm.neuro.uni-jena.de/vbm/). The ASL images were co-registered to the corresponding native grey matter images and spatially normalized to the MNI space using the deformation matrices from the T1 normalization step with a final isotropic resolution of 1.5 x 1.5 x 1.5 mm. The resulting ASL images were smoothed using an isotropic Gaussian filter of 10 mm. All images were visually inspected for proper realigning and spatial normalization.

### Statistical analyses

Three different analyses were performed. First, a whole brain voxel-by-voxel group analysis was performed comparing the smoothed and normalized ASL images from 12 children with ASD compared to 28 children with TD, using the framework of the general linear model within SPM8. Further region of interest (ROI) analyses was performed on the superior temporal regions, bilaterally generated with WFU PickAtlas software [27] and dilated 3 mm. Both analyses were performed on a grey matter mask thresholded to 50% and by scaling images proportionally to the individual global ASL signal. P values set to 0.05 Family Wise Error (FWE) corrected for multiple comparisons. In addition, a receiver operating characteristic (ROC) curve was generated with the rest CBF values from the cluster identified in the whole brain analysis (R statistical software *http://cran.r-project.org*). Secondly, an individual analysis was performed, in which the ASL image of each child with ASD was compared to the ASL images of the control group. Finally, whole brain SPM correlation analyses were performed to study univariate relationships between CBF and ADI-R score.

## Results

### Group Analysis

#### Whole brain voxel-by-voxel analysis

Whole brain voxel-by-voxel SPM group analysis revealed a significant hypoperfusion (t(38)=5.03, FWE corrected p<0.05) in the ASD group localized in the left superior temporal sulcus (MNI x, y, z coordinates: −60 −40 −2).

#### Regions of interest analysis

Analyses restricted to STG and STS revealed a significant decrease in rest CBF (p<0.05 corrected) in the ASD group compared to TD group bilaterally in the superior temporal sulcus (Fig 1A).

**Fig 1.**
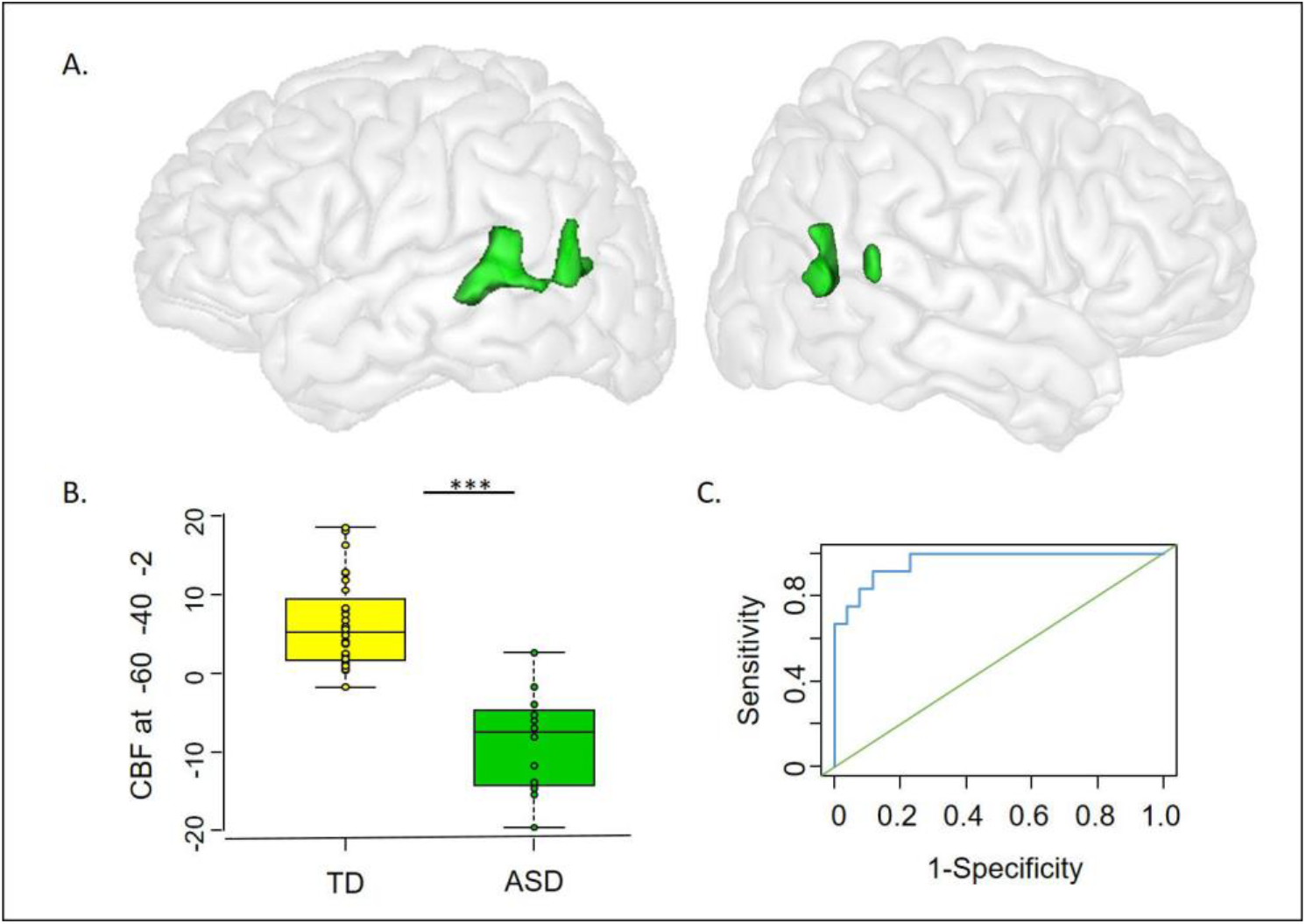
Group analysis revealed a significant rest CBF decrease (p<0.05, FWE corrected for multiple comparisons) in the ASD group compared to TD group bilaterally in the superior temporal sulci. (A) Regions with significant hypoperfusion (green), are superimposed on a rendering of the MNI-152 template average brain. (B) Box plot comparison of CBF signal between ASD and TD groups within the left superior temporal gyrus. (C). ROC curve analysis of rest CBF values within this region revealed an optimal cut-off value for which 91.7% of children with ASD in this sample were correctly identified as positive (91.7% sensibility) and 88.5% of TD children were correctly identified as negative (88.5% specificity).

#### Receiver operating characteristic (ROC)

The ROC curve analysis revealed an optimal cut-off rest CBF of 76.9 ml/100mg/min. Rest CBF values within the STS lower than the cut-off were observed in 11 out of 12 ASD patients (91.7% sensibility) while only 3 out of 26 TD children had lower than cut-off rest CBF values (88.5% specificity) (Fig 1B-C).

#### Individual Analysis

A significant temporal hypoperfusion (p<0.001 unc) was individually detected in 10 out of 12 children with ASD (83% of positive individual detection). Out of the 12 children with ASD, the temporal hypoperfusion was bilateral in 2 children, located on the right hemisphere in 4 children and on the left hemisphere in 4 children. Fig 2 shows brain regions with significant hypoperfusion in 3 children with ASD.

**Fig 2.**
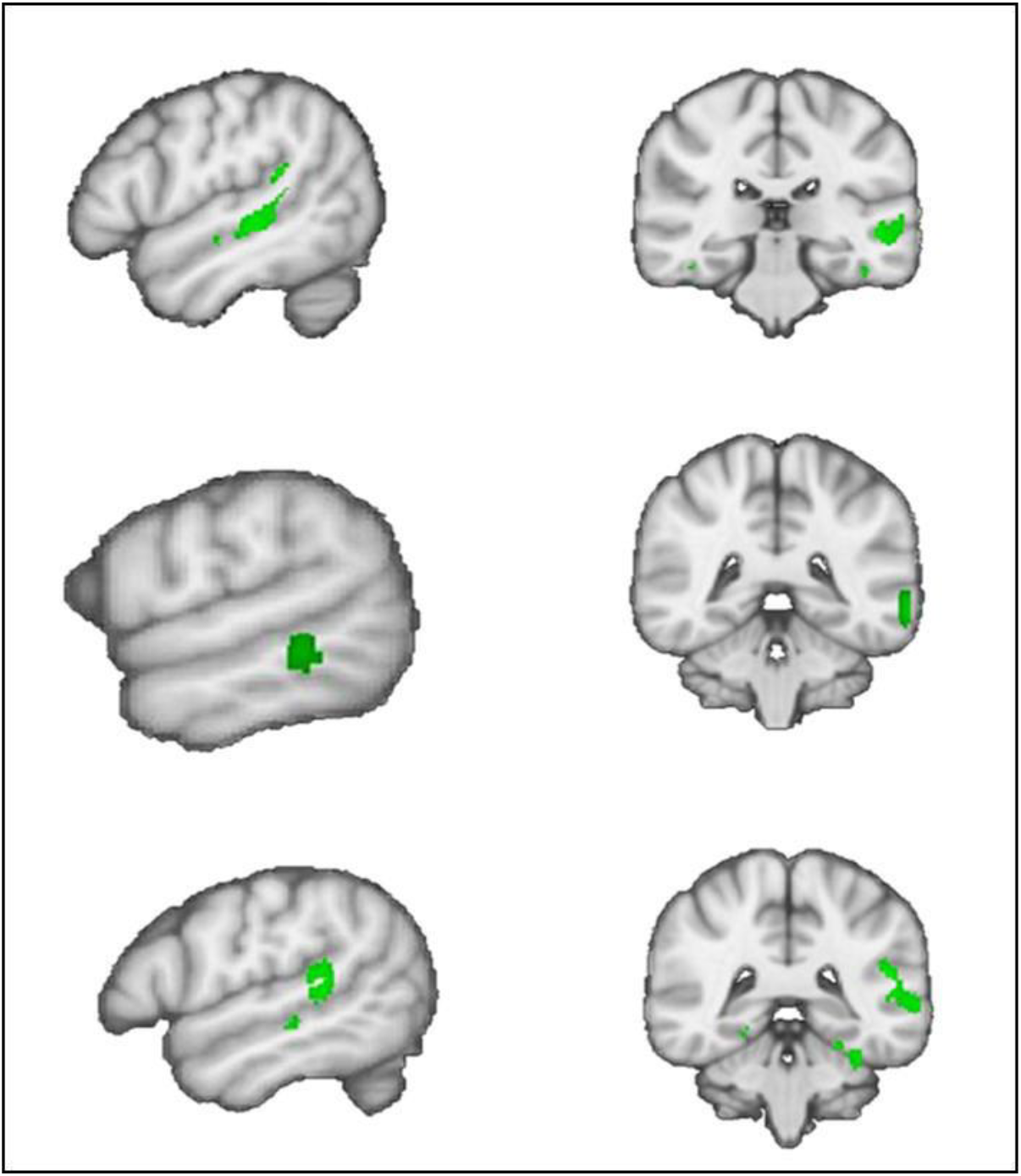
Examples of individual hypoperfusion in three children with ASD. Whole brain individual analysis revealed a significant temporal hypoperfusion (p<0.001 unc) in 10 out of 12 ASD patients (83% positive individual detection).

#### Correlation Analysis

Whole brain correlation analyses between rest CBF and ADI-R score showed significant negative correlation (p < 0.001, uncorrected; Talairach’s x, y, z coordinates: −50, −68, 24) located in the right superior temporal gyrus; more severe autistic symptoms are associated with lower CBF values in this region (Fig 3). In addition, a significant positive correlation between ADI-R score and rest CBF in the cerebellum (p < 0.001, uncorrected; Talairach’s x, y, z coordinates: −3, −52, 20); less severe autistic symptoms are associated with higher CBF values in this region.

**Fig 3.**
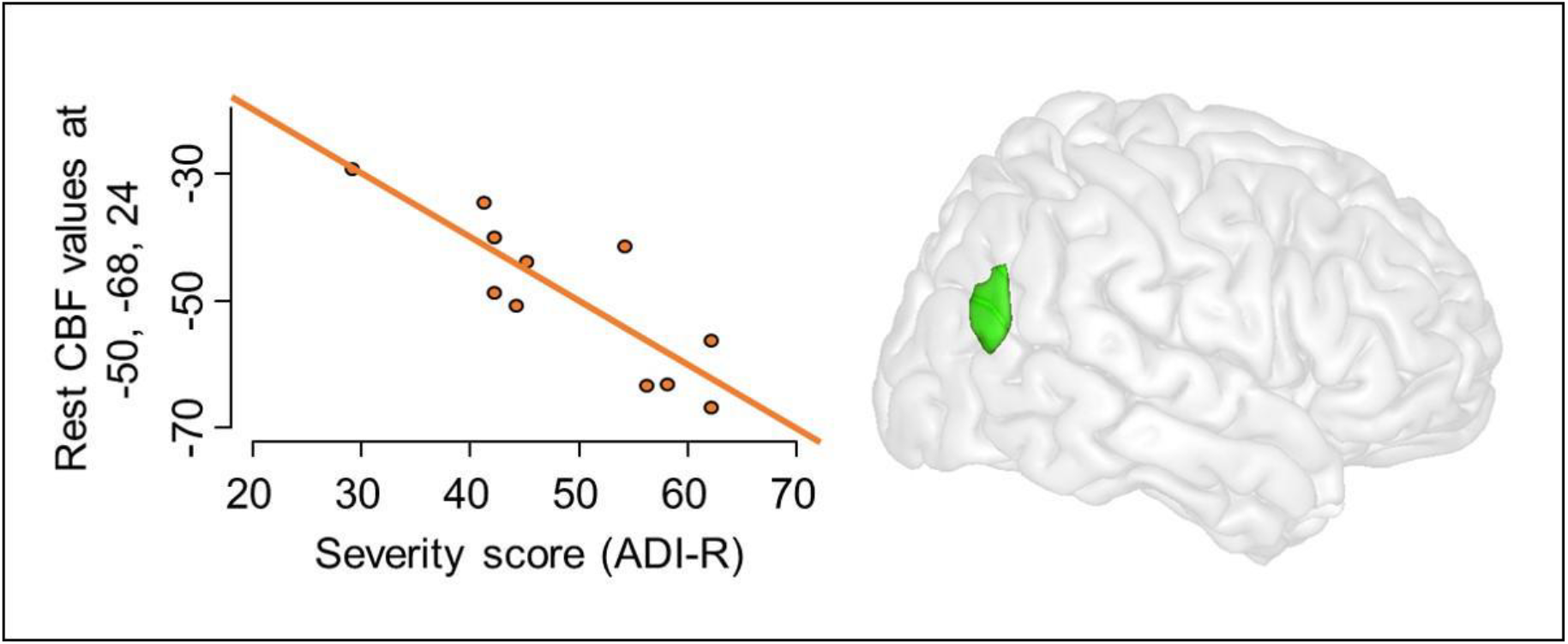
Plot from correlation analysis between ADI-R score and rest CBF values within the right pSTS (−50, −68, 24). p < 0.001 unc.

## Discussion

In the present study, using a new method allowing to investigate brain functioning at rest non-invasively using MRI, we report a significant decrease in rest CBF in children with ASD compared to typically developing (TD) children in superior temporal regions. Indeed, whole brain analysis revealed a significant decreased rest CBF in the left superior temporal sulcus (STS) and a further ROI analysis over both superior temporal regions confirmed a significant bilateral decrease is rest CBF in children with ASD. Moreover, ROC curve analysis of rest CBF values within this region revealed an optimal cut-off value for which 91.7% of children with ASD in this sample were correctly identified as positive (91.7% sensibility and 88.5% specificity). In addition, the temporal hypoperfusion was individually detected in 83% of the children with ASD. Finally, we found that ADI-R score, a global index of ASD severity, correlated negatively with rest CBF in the right superior temporal gyrus: children with more severe clinical scores, are those who have lower rest CBF values in this region, suggesting that right superior temporal hypoperfusion is related to ASD severity.

Strikingly, despite the small sample studied here, our results are extremely similar to previous PET and SPECT findings, which have shown rest hypoperfusion in the superior temporal regions in children with ASD when compared to control groups [10, 11]. Moreover, as in the present study, this hypoperfusion was also previously individually detected in a high percentage of children with ASD [11] and a significant negative correlation was observed between rest CBF and the ADI-R score in the superior temporal gyrus [12].

Brain imaging investigations on the neural basis of ASD raise the question of the pertinence of the results considering the whole wide autism spectrum. The first PET and SPECT studies on rest functional abnormalities in ASD describing decrease rest CBF in the temporal regions concerned children with ASD presenting intellectual disability. Here, we describe the same rest functional abnormality in children with ASD without intellectual disability. The congruence between these results in different ASD profiles reinforce the strength of rest functional abnormalities within this superior temporal regions in ASD and strongly indicates it’s is a core characteristic of the disorder.

Despite heterogeneity in results obtained in neuroimaging studies in ASD in the last two decades, anatomo-functional abnormalities within the temporal regions, particularly the STS, have been frequently described. Anatomical studies using voxel-based morphometry (VBM) showed reduced grey matter volume in the superior temporal regions [28–31]. Recently, a large database study described decreased cortical thickness in the temporal cortex [32]. Studies using diffusion tensor imaging (DTI) have indicate disrupted structure within white matter tracts in the brain, particularly in tracts connecting the temporal regions [33]. In addition, a large series of fMRI studies have described a lack of activation in the STS during socially relevant tasks, such as perceiving eye gaze as well as hearing human voice or performing socially relevant tasks [34–37].

Today, we know that the STS is highly implicated in processing social information, ranging from the perception of visual and auditory social-relevant stimuli (eye-gaze and voice) to the more complex processes of understanding the mental state of others (theory of mind) [38, 39]. Considering that, results showing rest functional abnormalities within the STS in ASD became relevant with regards to ASD core symptoms, reinforced here by the fact that the more severe the impairments are, stronger are abnormalities within the STS.

The results described here provide two important points for future investigations in ASD. Firstly, identifying a core dysfunctional region in ASD bears direct implications to the development of novel therapeutic interventions. For example, in depression disorder, classical rest functional abnormalities in frontal regions have been the ground to development of transcranial magnetic stimulation (TMS) protocols applied to this region, with significant clinical improvements [40, 41]. Results presented here suggest that STS could become an interesting target for this type of intervention in ASD. In a recent study, we have shown that, by disrupting the right pSTS neural network in healthy volunteers with inhibitory TMS, we artificially induced a gaze pattern that is similar to the gaze pattern observed in ASD, namely reduced eye-looking [42]. Difficulties in social perception processes, mainly characterized by reduced eye-looking as evidenced by eye-tracking studies, are a core symptom in ASD. If stimulation of the STS by excitatory TMS could change this pattern by inducing an increase in looking to the eyes, new perspectives on therapeutic interventions for ASD could emerge.

Secondly, multivariate classification analysis had shown that superior temporal rest hypoperfusion detected with PET allows prediction of ASD diagnosis with high accuracy, indicating it’s use as a potential biomarker in ASD [43]. However, methodological limitations such as radio isotopes injection prevented subsequent studies and its spread use. ASL-MRI allows to investigate rest CBF non-invasively and without such methodological limitations. Our results showing superior temporal rest CBF decrease in ASD with non-invasive ASL-MRI reinforce the robustness of this abnormalities. Searching for consensus on brain abnormalities in ASD, large-scale multi-site studies have been conducted over the past years using a multimodal neuroimaging approach. The present results suggest the potential interest of including ASL-MRI rest CBF measures in these studies. Confirmation of these results in a larger sample might indicate that hypoperfusion within the STS could truly become a biomarker in ASD.

